# Protein turnover dynamics suggest a diffusion to capture mechanism for peri-basal body recruitment and retention of intraflagellar transport proteins

**DOI:** 10.1101/2020.11.16.385518

**Authors:** Jaime V.K. Hibbard, Neftali Vazquez, Rohit Satija, John B. Wallingford

## Abstract

Intraflagellar transport (IFT) is essential for construction and maintenance of cilia. IFT proteins concentrate at the basal body, where they are thought to assemble into trains and bind cargoes for transport. To study the mechanisms of IFT recruitment to this peri-basal body pool, we quantified protein dynamics of eight IFT proteins, as well as five other basal body localizing proteins, using fluorescence recovery after photobleaching in vertebrate multiciliated cells. We found that members of the IFT-A and IFT-B protein complexes show distinct turnover kinetics from other basal body components. Additionally, known IFT sub-complexes displayed shared dynamics, and these dynamics were not altered during cilia regeneration as compared to homeostasis. Finally, we evaluated the mechanisms of basal body recruitment by depolymerizing cytosolic MTs, which suggested that IFT proteins are recruited to basal bodies through a diffusion-to-capture mechanism. Our survey of IFT protein dynamics provides new insights into IFT recruitment to basal bodies, a crucial step in ciliogenesis.

## INTRODUCTION

Construction and maintenance of cilia requires the movement of protein cargoes by a conserved active transport process termed intraflagellar transport (IFT) (Kozminski et al., 1993; Lechtreck, 2015; Sung and Leroux, 2013). IFT proteins act as adaptors between cargoes and the microtubule motors kinesin and dynein. Kinesin-II drives anterograde IFT, carrying cargoes from the cell body to cilium tip (Cole et al., 1998; Kozminski et al., 1995; Walther et al., 1994). Retrograde IFT is powered by IFT dynein and recycles signaling molecules and other proteins from the ciliary tip to the basal body at the base of cilia (Pazour et al., 1999; Pazour et al., 1998; Porter et al., 1999; Signor et al., 1999).

While IFT proteins have an essential axonemal function, the highest concentration of IFT proteins in the cell is found in a pool surrounding the basal body (Cole et al., 1998; Deane et al., 2001; Jurczyk et al., 2004; Lechtreck, 2015; Qin et al., 2004; Vashishtha et al., 1996). This peri-basal body pool is proposed to be the site of IFT cargo loading (Lechtreck, 2015), which is practical considering the higher local concentration of ciliary proteins within this pool can facilitate faster reaction kinetics. Additionally, perturbations that deplete the peri-basal body IFT pool are associated with loss of cilia (Čajánek and Nigg, 2014; Jurczyk et al., 2004; Richey and Qin, 2012; Toriyama et al., 2016), implicating the peri-basal body pool of IFT in the broad class of human diseases termed ciliopathies (Hildebrandt et al., 2011). Despite its importance for vertebrate development, the mechanisms by which IFT proteins are recruited to the peri-basal body pool from the cytoplasm remain unclear.

Multiple hypotheses could explain IFT protein localization to basal bodies. For example, IFT proteins might be actively transported to the basal body along cytosolic microtubules (MTs) (Fu et al., 2016; Hao et al., 2011; Mukhopadhyay et al., 2010). This hypothesis is enticing because the basal body serves as the microtubule organizing center for not only ciliary, but also cytosolic MTs (LeDizet and Piperno, 1986; Sandoz et al., 1988; Tucker, 1984). Indeed, ciliary protein Ccdc66 and ciliary receptor Rhodopsin are transported along cytoplasmic MTs via dynein motors (Conkar et al., 2019; Prosser and Pelletier, 2020; Tai et al., 1999). An alternative ‘diffusion to capture’ mechanism, whereby proteins freely diffuse until they find a docking site at their appropriate subcellular localization, plays an important role in localizing some ciliary proteins (Harris et al., 2016). Disruption of substructures of the basal body known as distal appendages, leads to loss of the peri-basal body IFT pool, so these structures may act as a capture site (Čajánek and Nigg, 2014; Deane et al., 2001; Schmidt et al., 2012; Singla et al., 2010; Yang et al., 2018).

Importantly, these mechanisms are not mutually exclusive, as the IFT proteins form complexes of multiple distinct subunits. These include two highly conserved protein complexes, IFT-A and IFT-B, composed of 6 and 16 proteins, respectively (Taschner and Lorentzen, 2016). IFT-A consists of both “core” and “peripheral” sub-units, and IFT-B is known to consist of two sub-complexes, IFT-B1 and IFT-B2 (Behal et al., 2012; Cole et al., 1998; Katoh et al., 2016; Lucker et al., 2005; Lucker et al., 2010; Mukhopadhyay et al., 2010; Piperno and Mead, 1997; Taschner et al., 2011; Taschner et al., 2016). At the basal body, these subunits, along with IFT motors, assemble into higher-order structures called trains (Webb et al., 2020). Whether subcomplexes share basal body recruitment and retention mechanisms remains unknown.

Recruitment and retention of IFT proteins near basal bodies are inherently dynamic processes, yet most results concerning the peri-basal body pool of IFT have been generated using static endpoint assays such as immunofluorescence. Recently, however, dynamic imaging has provided new insights into this important problem (Buisson et al., 2013; Mclnally et al., 2019; Wingfield et al., 2017; Yang et al., 2019). Notably, a recent study in *Chlamydomonas* has shown that the majority of the IFT peri-basal body pool is maintained via exhange with cytosolic IFT proteins (Wingfield et al., 2017). Furthermore, that study found that certain IFT-B members have similar kinetics of turnover in the peri-basal body pool, consistent with the idea that IFT-B subcomplexes are pre-assembled in the cytoplasm before recuitment to the basal body (Wingfield et al., 2017). However, a comprehensive survey of IFT protein dynamics at basal bodies has not been reported.

Finally, dynamic studies of the peri-basal body IFT pool have primarily been conducted in unicellular organisms such as *Chlamydomonas, Giardia,* and *Trypanosoma* (Buisson et al., 2013; Mclnally et al., 2019; Wingfield et al., 2017; Yang et al., 2019). Though many molecular mechanisms of ciliogenesis are shared between these organisms and vertebrates (Sigg et al., 2017; Sung and Leroux, 2013), other mechanisms, such as the CPLANE complex, are not evolutionarily conserved (Adler and Wallingford, 2017). We therefore quantified protein dynamics of several components of IFT-A and IFT-B, as well as other ciliary proteins at the basal body pool using fluorescence recovery after photobleaching (FRAP) in vertebrate multiciliated cells (MCCs). We discovered that members of the IFT-A and IFT-B protein complexes show distinct turnover kinetics, suggesting multiple mechanisms of IFT protein renewal in the basal body pool. Furthermore, we performed the first direct test of the hypothesis that cytosolic MTs could provide tracks for active transport of IFT proteins to basal bodies.

## RESULTS AND DISCUSSION

### Distinct turnover kinetics for ciliary proteins in the basal body pool

To quantify protein dynamics in vertebrate MCCs *in vivo,* we turned to the epidermis of *Xenopus laevis* embryos. This system has emerged in recent years as a key platform for *in vivo* analysis of IFT and other ciliary systems (Walentek and Quigley, 2017; Werner and Mitchell, 2012). We expressed fluorescently-tagged ciliary proteins using methods described previously (Brooks and Wallingford, 2015) and used confocal microscopy to image the *en face* apical optical section of MCCs, which contains basal bodies, and performed FRAP (Fig. 1A). We bleached small regions of interest (ROIs) containing three to five basal bodies within our field of view (Fig. 1B) and used a neighboring cell with non-bleached basal bodies outside the ROI to correct for changes to background fluorescence during imaging. Under our experimental conditions, halftimes were quite short (~20 sec for most proteins tested) (Supp. Fig. 1), which is consistent with a freely diffusing protein. We therefore used mobile fraction to indicate the fraction of basal body protein that was exchanging with a cytosolic pool versus stable in the basal body pool. On short timescales (seconds to minutes), proteins that are stably associated with the basal body will be less able to exchange protein contents and will have a lower mobile fraction value (Axelrod et al., 1976; Reits and Neefjes, 2001).

**Figure 1.**
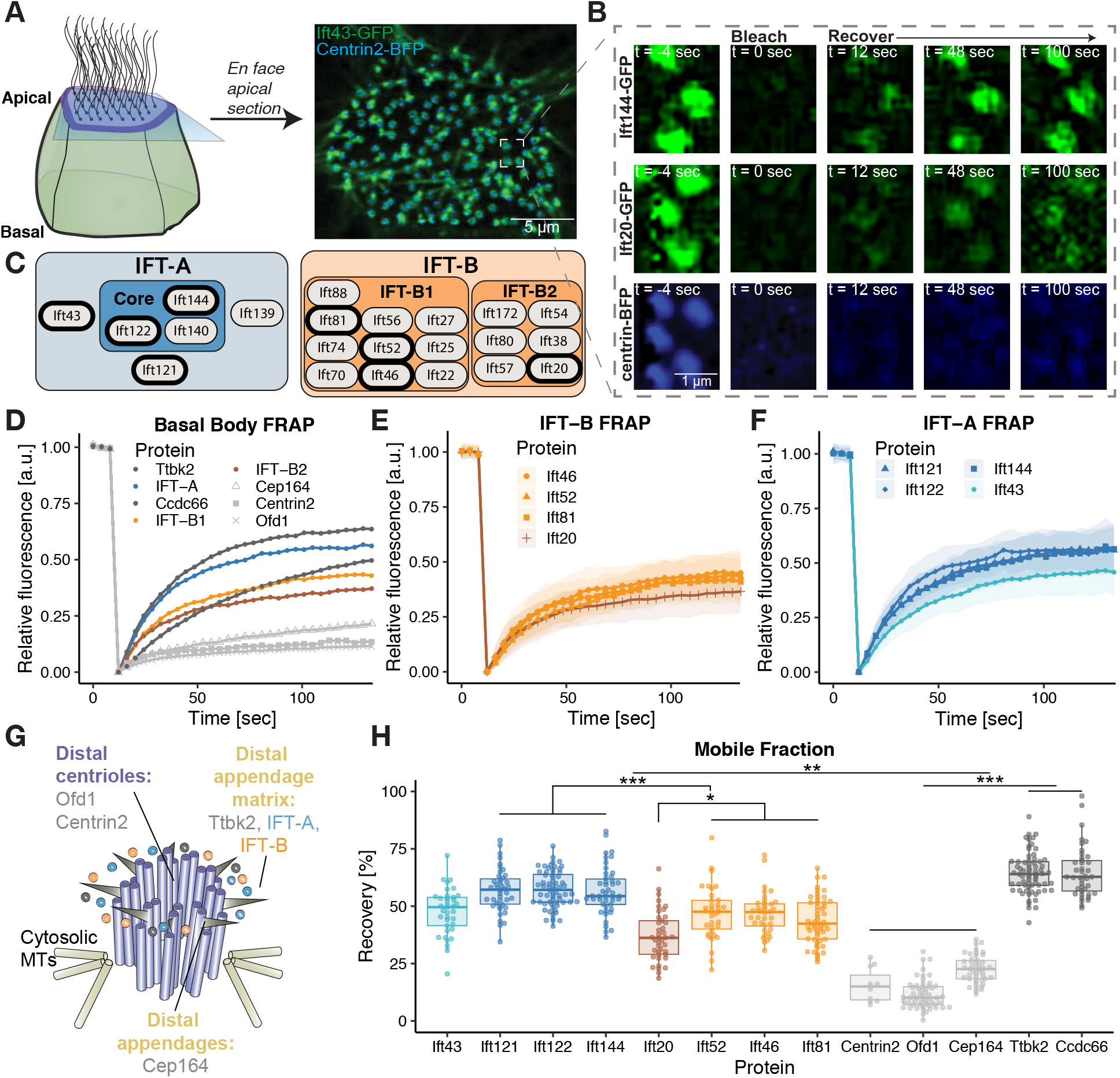
Functionally related proteins show distinct turnover kinetics in the basal body pool. (A) Schematic representation of FRAP experimental setup. Ift43-GFP accumulates around Centrin2-BFP, a marker for basal bodies. (B) Representative images from FRAP experiments of Ift144-GFP, Ift20-GFP, and Centrin2-BFP. (C) Schematic of IFT-A and IFT-B complexes. Proteins investigated in this report are bolded. (D) FRAP recovery curves of IFT-A proteins (averaged from Ift144-GFP, GFP-Ift122, and Ift121-GFP), IFT-B1 proteins (averaged from Ift81-GFP, Ift52-mNG, and Ift46-mNG), IFT-B2 protein Ift20-GFP, structural basal body components (Cep164-GFP, Centrin2-BFP, GFP-Ofd1), and basal body regulators (GFP-Ttbk2, GFP-Ccdc66). For clarity of presentation, error bars are removed. (E) FRAP recovery curves of IFT-B proteins. Shaded area corresponds to standard deviation. (F) FRAP recovery curves of IFT-A proteins. Shaded area corresponds to standard deviation. (G) Schematic representation of a basal body, showing the localization of different proteins. (H) Mobile fraction quantification of basal body turnover kinetics for investigated proteins. Several statistical differences are noted in the tree, for full discussion of differences, see Supp. Table 2.

We first established baseline parameters by performing FRAP on a diverse group of known basal body-associated proteins (Fig. 1G). As expected for structural components of the basal body, we found that BFP or GFP fusions to Centrin2, Ofd1, and Cep164 displayed very small mobile fraction values (Fig. 1B, D, H), indicating that they are very stably retained and exchange slowly with the cytoplasm (Paoletti et al., 1996; Schmidt et al., 2012; Singla et al., 2010). By contrast, known ciliogenesis regulator Ttbk2 (Goetz et al., 2012), displayed a much higher mobile fraction, suggesting it is not stably retained but rather exchanges more freely with the cytoplasm. Finally, Ccdc66 is implicated in the recruitment of ciliary proteins and displayed an intermediate FRAP profile (Conkar et al., 2019; Conkar et al., 2017), far more dynamic than the structural components, but slower than Ttbk2. These data demonstrate the efficacy of our experimental platform for exploring protein dynamics and suggest that ciliary and centriolar proteins display diverse patterns of turnover at basal bodies in MCCs.

### Functionally related IFT subcomplexes display shared protein dynamics

We next sought to use our FRAP platform to explore the dynamics of IFT protein recruitment and retention in the peri-basal body pool. We chose eight IFT proteins (bold outlines in Fig. 1C) for analysis to sample the dynamics of the full range of known IFT sub-complexes.

The first trend that was clear in our data was that known sub-complexes displayed shared dynamics. For example, the IFT-B complex is known to be composed of two sub-complexes, IFT-B1 and IFT-B2 (Taschner et al., 2016). We examined the kinetics of recovery for three sub-units of the IFT-B1 complex. Ift46, Ift52, and Ift81, and all showed similar mean mobile fraction values (47%, 47%, and 43%, respectively). Strikingly, however, the IFT-B2 protein Ift20 displayed significantly different dynamics, with a mean mobile fraction value of 37% (Fig. 1E, H). The distinct turnover kinetics suggests the possibility that the IFT-B1 and IFT-B2 subcomplexes are retained in the peri-basal body pool by distinct mechanisms.

We also observed differences in the turnover dynamics of IFT-A proteins. First, all IFT-A components tested displayed significantly greater mobile fractions as compared to IFT-B (Fig. 1H). IFT-A is comprised of biochemically distinct “core” and “peripheral” subunits (Behal et al., 2012; Mukhopadhyay et al., 2010; Zhu et al., 2017). Two core (Ift122/Ift44) and one peripheral protein (Ift121) all displayed shared dynamics, each with a mean mobile fraction value of ~57% (Fig. 1H). Finally, the peripheral IFT-A component Ift43 consistently displayed different turnover dynamics when compared to the other IFT-A proteins (Fig. 1F, H). Importantly, this finding is consistent with multiple studies suggesting Ift43 is sub-stoichiometric in the IFT-A complex (Behal et al., 2012; Mukhopadhyay et al., 2010).

These data in vertebrate MCCs confirm and extend previous findings in *Chlamydomonas* (Wingfield et al., 2017) and establish shared basal body turnover dynamics within functionally related IFT subcomplexes as an evolutionarily conserved feature. Moreover, these conserved turnover dynamics suggest that IFT protein complexes are likely recruited and retained at basal bodies as assembled subcomplexes, rather than as individual subunits. Coupled to biochemical findings that many individual IFT proteins are unstable without their binding partners (Bhogaraju et al., 2013; Richey and Qin, 2012; Taschner et al., 2014), our live imaging data suggests that IFT subcomplex assembly likely occurs in the cytoplasm, rather than in the basal body pool.

### Intra-IFT protein interactions control IFT basal body turnover kinetics

To explore the significance of the observed pattern of IFT protein dynamics in the peri-basal body pool, we next asked if known protein-protein interactions within IFT complexes impacts IFT protein turnover. We focused on Ift46, an IFT-B1 member, because it is known to be recruited to the basal body by its interaction with Ift52 in *Chlamydomonas* (Lv et al., 2017). First, we confirmed that this functional interaction is conserved in *Xenopus* by expressing Ift46 that lacked its C-terminal Ift52 binding domain (Ift46ΔC-mNG). We observe that while full-length Ift46-mNG was properly recruited to the basal body, Ift46ΔC-mNG was not and instead displayed strong diffuse signal throughout the cytoplasm (Fig. 2A). This result further validates the conservation of basal body recruitment mechanisms between *Chlamydomonas* and vertebrates.

**Figure 2.**
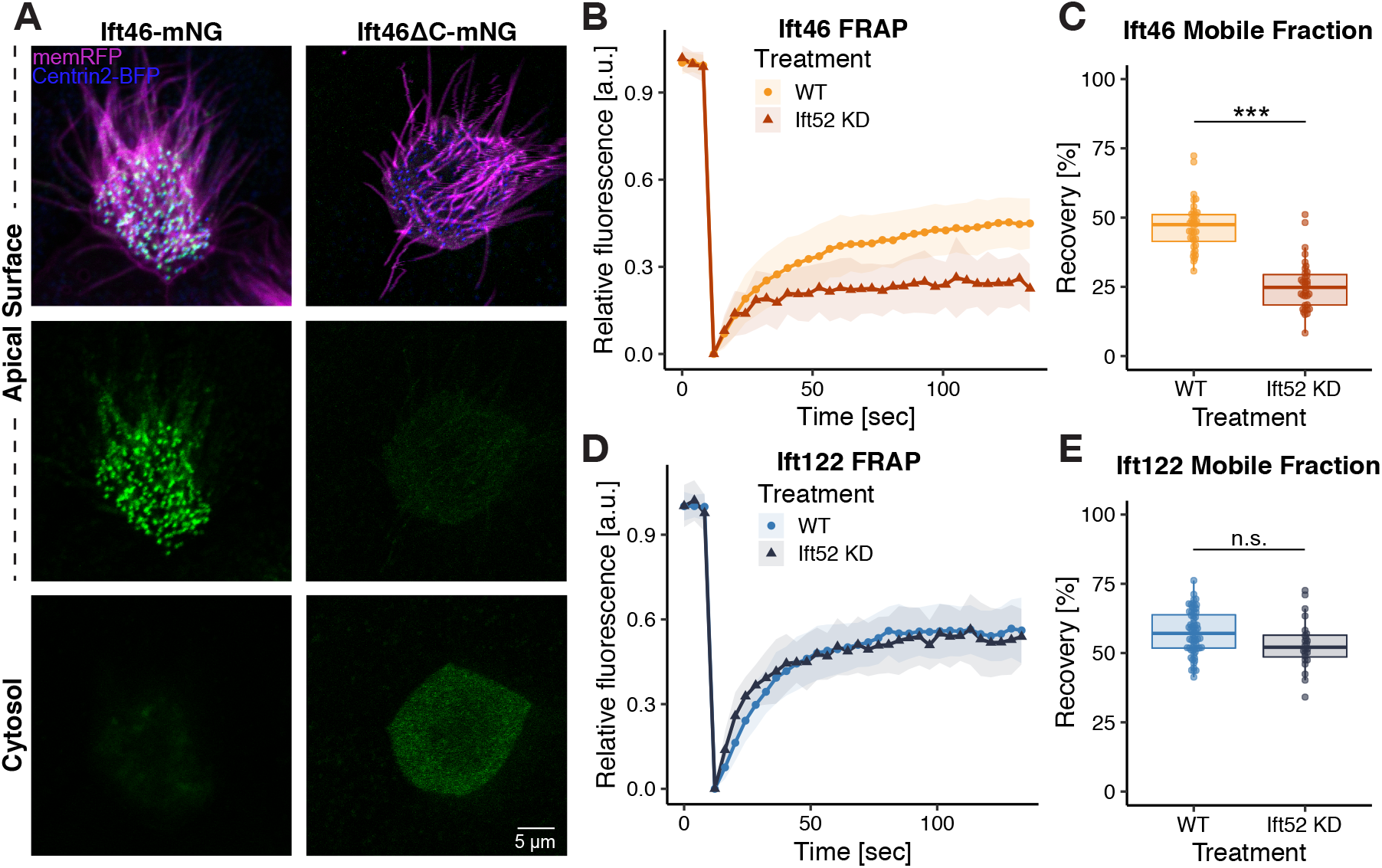
Ift52 is required for the recruitment and normal turnover of Ift46 at basal bodies. (A) Imaging of Ift46-mNG (left panel) and Ift46ΔC-mNG (right panel). (B) FRAP of WT Ift46-mNG, compared to *ift52* KD. (C) The mobile fraction value of Ift46-mNG is significantly lower upon KD of Ift52. (D) FRAP recovery of GFP-Ift122, an IFT-A protein, is not affected by *ift52* KD. (E) Mobile fraction quantification of GFP-Ift122 in WT and *ift52* KD embryos.

As Ift46 recruitment to basal bodies is dependent on its interaction with Ift52 (Lv et al., 2017), we hypothesized that the disruption of Ift52 would lead to changes in basal body turnover dynamics of Ift46, so we assessed FRAP turnover after knock-down (KD) of *ift52* with a previously described morpholino (Dammermann et al., 2009; Drew et al., 2017). KD of *ift52* leads to decreased levels of Ift46-mNG at basal bodies, as expected, and importantly, this phenotype could be rescued by re-expression of Ift52 (Supp. Fig. 2). Consistent with the hypothesis that Ift52 loss will disrupt the FRAP kinetics of its binding partner, we determined that Ift46 displayed significantly reduced turnover in the basal body pool upon KD of *ift52* (Fig. 2B, C). As there is still remnant Ift46 enrichment at the basal body (Supp. Fig. 2), we hypothesize that *ift52* KD reduces the number of basal body binding spots available to Ift46, resulting in less dynamic turnover.

As a control for specificity, we noted that IFT-B is not required for IFT-A recruitment to basal bodies (Brown et al., 2015), and we found that Ift122 dynamics were unaffected by *ift52* KD (Fig. 2D, E). These data validate the utility of our FRAP system to explore the mechanisms of IFT retention at basal bodies.

### IFT recruitment kinetics at basal bodies are unaltered during cilia regeneration

The IFT retention kinetics described above were obtained in full-length cilia, which homeostatically maintain a relatively constant length. Interestingly, during cilia regeneration, several aspects of IFT are altered, as substantially more cargoes must be transported to build the growing axoneme. Indeed, flagellar regeneration in *Chlamydomonas* is characterized by changes in the size of the peri-basal body pool, IFT train size, and in cargo loading onto IFT trains (Craft et al., 2015; Engel et al., 2009; Ludington et al., 2013; Wren et al., 2013). On the one hand, ciliary growth requires the import of thousands of copies of ciliary precursors (Qin et al., 2004), so we might expect to observe altered kinetics of IFT during regeneration. Alternatively, steady-state basal body turnover of IFT could be sufficient to ‘prime’ cilia for regeneration. In this model, an increase in cargoes per IFT train could be sufficient to increase ciliary cargo import, without the need to change IFT recruitment to basal bodies. Consistent with this latter hypothesis, cargo-loading on IFT trains scales inversely with ciliary length (Craft et al., 2015; Engel et al., 2009; Wren et al., 2013).

To ask if IFT turnover rates differ during ciliary growth, we used a previously described method for deciliation in *Xenopus* MCCs (Werner and Mitchell, 2013) and then quantified the time-course of cilia regeneration using confocal imaging at 30 min intervals. *Xenopus* MCCs underwent an obvious lag phase before beginning regrowth, with no evident increase in cilia length in the first 60 min (Fig. 3A, B). Cilia then grew at a fairly consistent rate over the next two hours (Fig. 3B).

**Figure 3.**
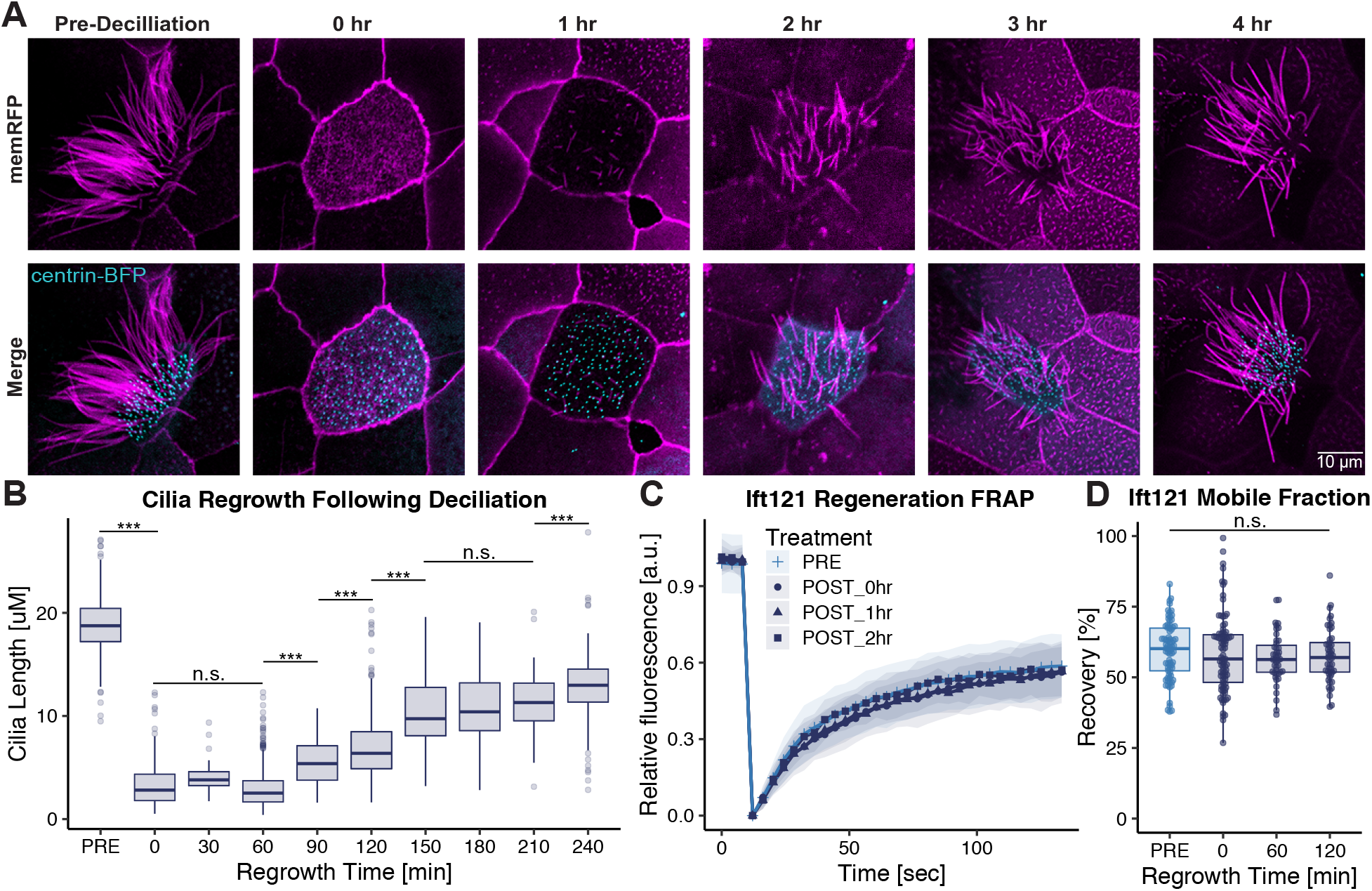
Induced cilia regeneration has no effect on IFT protein dynamics at the basal body. (A) Time course tracking cilia regrowth, pre deciliation, post deciliation and every hour up to the fourth hour timepoint. (B) Quantification of ciliary length prior to and during ciliary regeneration. (C) FRAP recovery curves of Ift121-GFP protein at different timepoints pre, post, and during ciliary regeneration. (D) Mobile fraction values of IFT121-GFP protein pre deciliation and at one and two-hour timepoints during ciliary regeneration.

We then examined basal body turnover of the IFT-A protein Ift121 immediately following deciliation and at early and mid-regrowth timepoints (Fig. 3C). Interestingly, we observed no change in the basal body turnover of Ift121 during cilia regeneration (Fig. 3D). These data suggest that the recruitment of Ift121 to the basal body is not adjusted during ciliary regeneration. Based on our previous data, we expect other members of the IFT-A complex to share basal body recruitment mechanisms. We therefore expect steady state turnover of IFT-A to be sufficient to ‘prime’ cilia for regeneration.

Additionally, this data further demonstrates that the basal body pool of Ift121 is maintained by exchange with the cytosolic pool, not through recycling from the cilia. This is consistent with data that suggest an “open” system for IFT-A, where IFT-A proteins returning from the axoneme following retrograde transport immediately re-enter the cytosolic pool (Wingfield et al., 2017).

### Cytosolic microtubules are dispensable for IFT recruitment to basal bodies

Next, we sought to explore the mechanisms of IFT transport to basal bodies from the cytosol. To investigate the potential that IFT proteins are themselves a cargo for microtubule-based active transport to basal bodies, we developed a system to specifically depolymerize cytosolic microtubules in *Xenopus* MCCs *in vivo.* Cold-shock techniques are known to depolymerize cytosolic microtubules without affecting ciliary microtubules (Behnke and Forer, 1967; Burton, 1968). However, cytosolic microtubules quickly reform upon return to room temperature. Because nocodazole inhibits microtubule polymerization (De Brabander et al., 1976; Hoebeke et al., 1976), we used a combination treatment of cold shock and nocodazole (CS+noc). Transverse sections of *Xenopus* MCCs (Fig. 4A, B) show this treatment effectively depolymerized cytosolic microtubules (Fig. 4C), without affecting ciliary microtubules (Fig. 4D).

**Figure 4.**
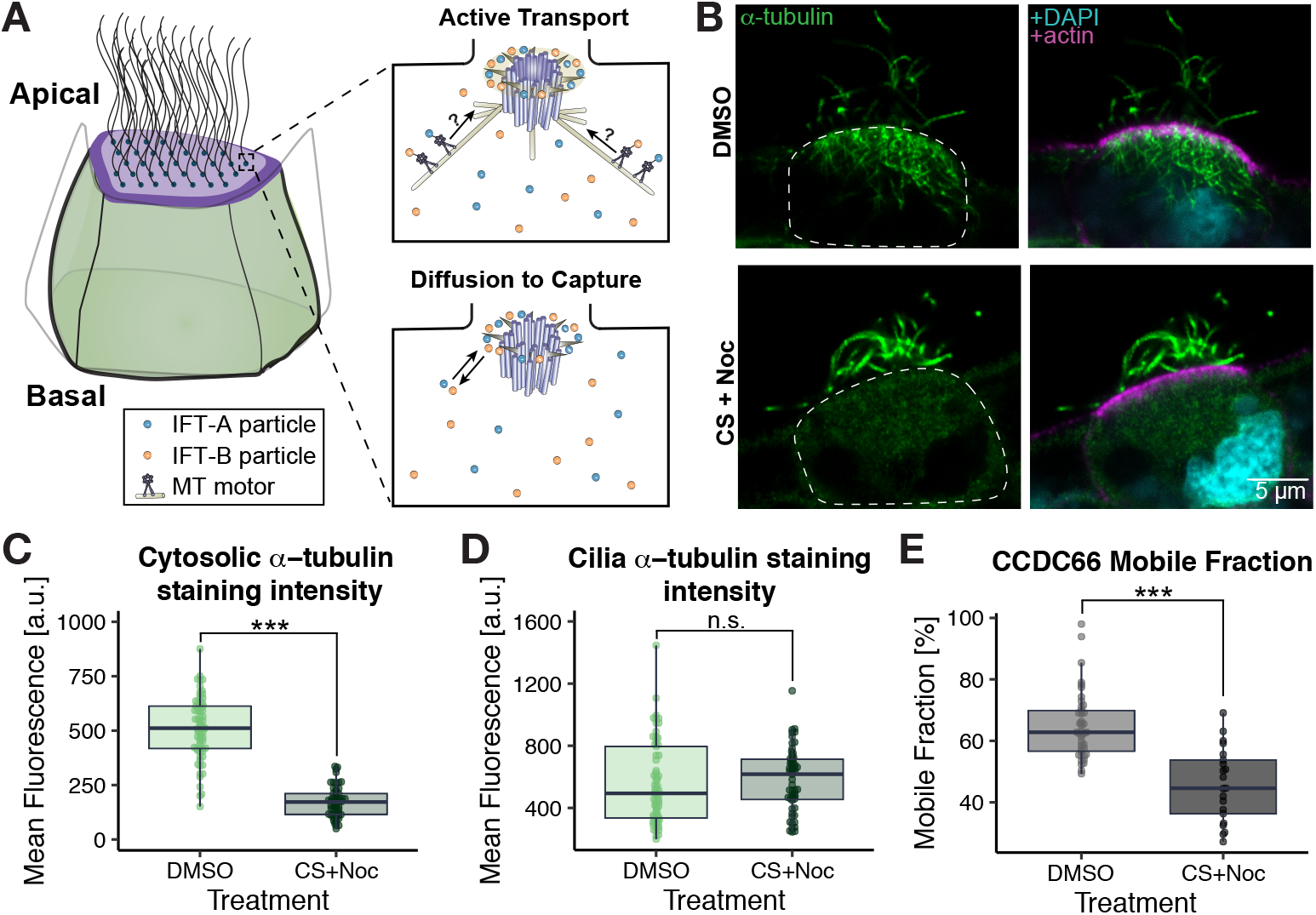
Cold-shock + nocodazole (CS + Noc) treatment eliminates cytosolic microtubules and disrupts the dynamics of Ccdc66 at basal bodies. (A) Schematic representation of MCC, depicting the two proposed mechanisms for IFT recruitment to basal bodies: active transport (upper panel) and diffusion to capture (lower panel). (B) Transverse sections of *Xenopus* MCCs stained for a-tubulin to verify efficacy of treatment. Actin (stained by Alexa Fluor 555 Phalloidin) and DAPI show the apical domain and nuclei, respectively. In DMSO (mock)-treated cells are shown in the upper panels and CS+Noc treated cells in the lower panels. (C) Cytosolic a-tubulin staining quantification for DMSO versus CS+Noc treated embryos. (D) Ciliary a-tubulin staining quantification for DMSO versus CS+Noc treated embryos. (E) Mobile fraction comparison of basal body turnover kinetics of Ccdc66 between DMSO and CS+Noc treatment.

To confirm this approach disrupted active transport to basal bodies, we examined the turnover kinetics of Ccdc66, which is known to require cytosolic microtubules for its localization to the base of primary cilia (Conkar et al., 2019). FRAP following CS+noc treatment of MCCs revealed a significant decrease in the mobile fraction value for Ccdc66, as compared to the untreated control (Fig. 4E). The ~19% mobile fraction decrease is consistent with the ~15% decrease observed for centrosomal localization in RPE cells (Conkar et al., 2019).

By contrast, all tested IFT proteins displayed no change after CS+noc treatment. FRAP recovery kinetics of IFT-A proteins Ift121, Ift122, and Ift144 appeared identical with and without CS+noc treatment (Fig. 5A). Similarly, FRAP kinetics of IFT-B proteins Ift52 and Ift20 did not change with CS+noc treatment (Fig. 5B). There was no statically significant change in the mobile fraction values of any IFT protein after disruption of cytosolic microtubules (Fig. 5C, D). These data demonstrate that cytosolic microtubules are dispensable for normal retention of IFT proteins at basal bodies, suggesting that a diffusion to capture mechanism is responsible for maintenance of the IFT basal body pool.

**Figure 5.**
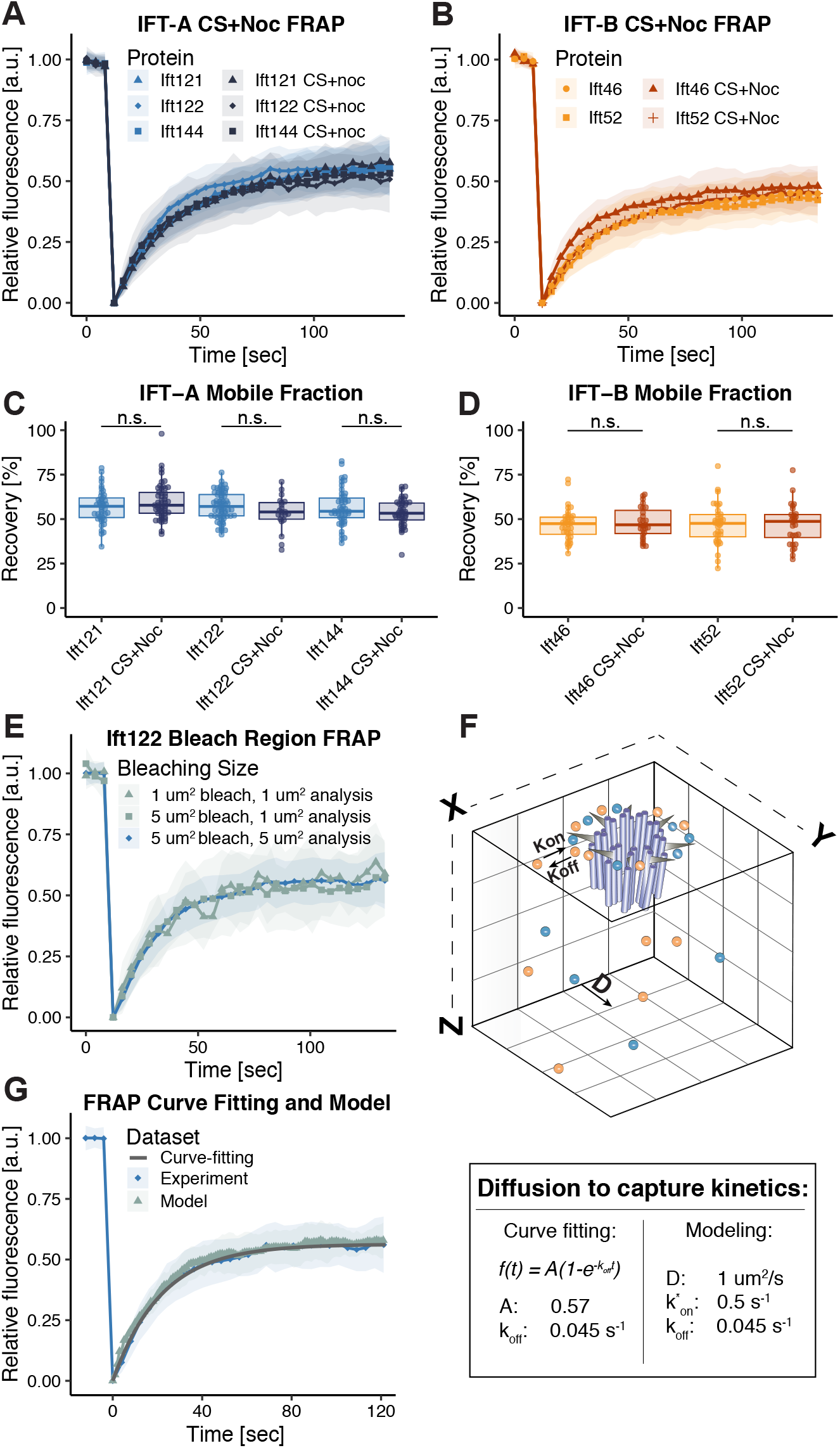
Cytosolic microtubules are dispensable for IFT recruitment to basal bodies. (A) FRAP recovery curves of IFT-A proteins with and without CS+Noc treatment. (B) FRAP recovery curves of IFT-B proteins with and without CS+Noc treatment. (C) Mobile fraction comparison of basal body turnover kinetics of IFT-A proteins between no treatment and CS+Noc treatment. (D) Mobile fraction comparison of basal body turnover kinetics of IFT-B proteins between no treatment and CS+Noc treatment. (E) FRAP recovery curves of GFP-Ift122 with 5 μm^2^ and 1 μm^2^ bleaching regions. (F) Experimental and model FRAP recovery curves of GFP-Ift122, with the curve-fitting equation line overlaid.

### *In vivo* quantification of IFT protein diffusion and binding kinetics by analysis and modeling of FRAP data

Elucidation of kinetic rate constants is essential for complete understanding of biological processes. To determine the kinetic rate constants of the diffusion to capture mechanism for IFT transport to basal bodies, we used a combination of curve fitting and modeling techniques.

When FRAP recovery is dominated by binding to a subcellular structure, FRAP curvefitting allows for elucidation of binding kinetics (Sprague and McNally, 2005). Therefore, we first verified that our FRAP recoveries were within a binding-dominant regime. To do so, we changed the size of our bleaching and analysis region from 5 μm^2^ to 1 μm^2^. If diffusion is a major contributor to FRAP recovery, smaller bleaching and analysis regions, which result in shorter travel distance for non-bleached fluorescent proteins, should demonstrate faster recoveries (Sprague et al., 2004). Performing this experiment with with IFT-A protein, Ift122, we found that FRAP recovery kinetics were identical between the smaller and larger bleaching regions, indicating that our recovery is in a binding-dominant regime (Fig. 5E).

Having established a binding-dominant regime, we then utilized curve-fitting to quantify the off rate for basal body binding (*k_off_*) and the mobile fraction *(A)* (Sprague et al., 2004). Fluorescence recovery as a function of time f(t) was fit to the following equation: *f*(*t*) = *A*(1 – *e^-k^off^t^*) and determined an off-rate of 0.045 s^-1^ (Fig. 5G).

Next, we sought to further characterize the kinetics of this diffusion to capture method by using a kinetic modeling approach (Phair and Misteli, 2001). We used a Monte Carlo model to simulate IFT particles moving on a lattice, with a diffusion coefficient *D* (Fig. 5F). If these particles encountered the basal body, simulated by a capture site with a boundary of 400×400 nm, they displayed a probability of binding dependent on the pseudo-first-order rate constant for binding (*k*_on_*) (Sprague et al., 2004). Molecules that were captured displayed a probability of unbinding dependent on the previously established *k_off_* = 0.045 s^-1^. Using this method, we simulated the turnover FRAP experiments and determined that D = 1 μm^2^/s and *k*_on_* = 0.5 s^-1^ closely recapitulated experimental data for representative IFT-A protein GFP-Ift122 (Fig. 5G). As expected for a protein of >100 kDa that is a member of a ~1 MDa complex, this diffusion coefficient is slower than cytosolic GFP (Potma et al., 2001), but faster than mRNAs undergoing translation (Yan et al., 2016).

## Conclusions

Elucidating the mechanisms of IFT proteins’ dynamic localization from the cytoplasm to the peri-basal body pool and their retention there is a crucial challenge in understanding both ciliogenesis and cilia homeostasis. While it has become evident that the distal appendage matrix appears to be the docking site for IFT proteins in the peri-basal body pool (Deane et al., 2001; Yang et al., 2018), the specific capture mechanism is still unclear. *In vitro* reconstitution of the centrosome, which is a modified centriole like basal bodies, elucidated the molecular requirements for microtubule nucleation (Stearns and Kirschner, 1994; Woodruff et al., 2017). *In vitro* reconstitution was similarly essential for elucidating the architecture of the IFT-B complex (Taschner et al., 2016). The quantification of *in vivo* rate constants presented here will provide an important foundation for future *in vitro* efforts to reconstitute IFT binding to the distal appendage matrix. Additionally, it will be interesting to further elucidate the changes to these rate constants using perturbations that have been uncovered to affect the peri-basal body pool of IFT with static endpoint assays such as immunofluorescence (Čajánek and Nigg, 2014; Schmidt et al., 2012; Singla et al., 2010). Finally, it will be interesting to determine how peri-basal body capture mechanisms influence the assembly of IFT machinery into trains.

## ACKNOWLEDGEMENTS

The authors thank D. Dickinson, E. Roberson, and N. Stolpner for critical discussions and reading. Work in the JBW Lab was supported by the NICHD (R01HD085901), NHLBI (R01HL117164). JVKH was supported by the Provost’s Graduate Excellence Fellowship from UT Austin and NV was supported by the Dean’s Strategic Fellowship from UT Austin. RS was supported by NSF grant CHE 1566001.

## MATERIALS AND METHODS

### *Xenopus* embryo manipulations

All experiments were conducted following animal ethics guidelines of the University of Texas at Austin, protocol number AUP-2018-00225. Female adult *Xenopus laevis* were induced to ovulate by injection of human chorionic gonadotropin. The following day, females were squeezed, and eggs were fertilized *in vitro* with homogenized testis. Embryos were dejellied in 1/3X Marc’s modified Ringer’s (MMR) with 3% (w/v) cysteine (pH 7.9) at the two-cell stage. All embryos were reared in 1/3X MMR solution until the appropriate stage. For microinjections, embryos were placed in 2% Ficoll in 1/3X MMR and injected using a glass needle, forceps, and an Oxford universal micromanipulator.

### Immunostaining

Immunostaining protocol was modified from (Lee et al., 2008). Stage 27 *Xenopus* embryos were fixed using 1X MEMFA for 2h on a rotating mixer at room temperature immediately following either DMSO or cold-shock and nocodazole treatment. For transverse sections, embryos were embedded in 2% agarose, and thick (~250 μm) sections were cut using a Vibratome 1000 system. Sections were further fixed in 1X MEMFA for 1h at room temperature.

Both whole-mount embryos and transverse sections were washed in TBST (155 mM NaCl, 10 mM Tris-Cl, 0.1% Triton X-100, pH 7.4) and blocked in 10% NGS, 5% DMSO in TBST. Monoclonal mouse anti-alpha-tubulin (DSHB, AA4.3, 1:200 dilution) was used as a primary antibody. Anti-alpha-tubulin was detected using Alexa Fluor 488-goat anti mouse (Invitrogen, 1:1000). In addition, nuclei were stained using DAPI (1:1000) and actin was stained using Alexa Fluor 568 Phalloidin (Invitrogen, 1:500). Images were acquired a Zeiss LSM700 laser scanning confocal microscope using a 100X oil objective lens.

Ciliary and cytosolic alpha-tubulin staining intensity quantification was performed by drawing an ROI around the ciliary or cytosolic regions of a single MCC and measuring the mean intensity of that region using Fiji (Schindelin et al., 2012). For comparisons of alpha-tubulin intensity between DMSO and CS+noc treated embryos, a minimum of 55 cells from 4 embryos were quantified per condition.

### Plasmids and cloning

Previously described vectors containing the following inserters were used: Ift43-GFP (Brooks and Wallingford, 2013), Ift121-GFP (Toriyama et al., 2016), GFP-Ift122 (Toriyama et al., 2016), Ift144-GFP (Toriyama et al., 2016), Ift20-GFP (Brooks and Wallingford, 2012), Ift81-GFP (Toriyama et al., 2016), GFP-Ofd1 (Toriyama et al., 2016), Cep164-GFP (Toriyama et al., 2016), GFP-Ttbk2 (Tu et al., 2018), Centrin2-BFP (Tu et al., 2018), and a-tub MAP7-RFP.

For new inserts, gene sequences were obtained from Xenbase (www.xenbase.org) (Karimi et al., 2018). Open reading frames were amplified from a *Xenopus* cDNA library by polymerase chain reaction and then inserted into either a pCS10R or pCS vector. All constructs were verified by sequencing.

### mRNA, morpholino, and plasmid microinjections

Vectors were linearized by restriction digestion and capped mRNAs were synthesized using the mMESSAGE mMACHINE SP6 transcription kit (ThermoFisher Scientific, #AM1340).

For knock-down experiments, we used the previously described antisense morpholino oligonucleotides (MOs) against *Ift52,* 5’-AAGCAATC TGTTTGTTGACTCCCAT-3’ (GeneTools) (Dammermann et al., 2009; Drew et al., 2017).

*Xenopus* embryos were injected with mRNAs, plasmids and/or morpholino oligonucleotides in the two ventral blastomeres at the four-cell stage to target the epidermis. mRNAs were injected at 70-200 pg per blastomere, Ift52 morpholino was injected at 40 ng per blastomere, and plasmids were injected at 25-75 pg per blastomere.

### Live imaging, FRAP, and image analysis

For live imaging, *Xenopus* embryos (stage 26-29) were mounted between two coverslips and submerged in 0.3X MMR and imaged using a Nikon A1R scanning confocal microscope with a 60X oil immersion objective.

For FRAP experiments, a region of interest (ROI) of either 2.5×2.0 μm (all FRAP experiments unless otherwise stated) or 1×1 μm, focusing on the basal bodies of a MCC, was defined. Bleaching was performed on the ROI using 35% laser power of either a 405 or 488-nm wavelength laser. Images of the bleached cell and a neighboring non-bleached MCC (which was used for subsequent bleach correction) were acquired. Bleach correction and FRAP curve-fitting was carried out using a custom python script (modified from http://imagej.net/Analyze_FRAP_movies_with_a_Jython_script). For WT FRAP analysis of each IFT protein, a minimum of 35 cells from at least 5 independent embryos in three different clutches were analyzed. For complete numbers, see Supp. Table 1.

The fluorescence intensity of Ift46-mNG at basal bodies was measured as previously described (Toriyama et al., 2016). Briefly, basal bodies were determined using the three-dimensional object counter plugin of Fiji software, with the object size filter minimum set to 20 and the threshold set to the plugin’s recommended value. At least 12 basal bodies per cell from at least 24 cells in 3 independent embryos were analyzed.

### Cold shock and drug treatment

To depolymerize cytosolic microtubules, a combination of cold shock and nocodazole treatment (CS + noc) was used. First, embryos were equilibrated in a solution of 25 μM nocodazole in 0.1X MMR for a minimum of 15 min. Cold-shock was performed by adding the embryos to 25 μM nocodazole in 0.1X MMR pre-chilled to 5°C for 15 min. Embryos were allowed to return to room temperature and maintained in 25 μM nocodazole in 0.1X MMR for imaging.

To ensure efficient and persistent depolymerization of microtubules, treated embryos utilized for FRAP live imaging were co-injected with the microtubule binding domain ensconsin of MAP7 (α-tubulin MAP7-RFP). Non-treated embryos showed robust ensconsin labeling of cytosolic microtubules, while CS+noc treated embryos only showed ensconsin signal at basal bodies. For FRAP analysis of CS+noc treated proteins, a minimum of 20 cells from 4 independent embryos was utilized.

### Deciliation and ciliary regeneration

Deciliation was performed as previously described (Werner and Mitchell, 2013). Briefly, embryos were incubated in deciliation buffer (75 mM calcium, 0.02% NP40 in 0.1X MMR+antibiotic (50 μg/mL gentamicin)) at room temperature for 5min. Embryos were washed in 0.1X MMR + antibiotic for 2min, maintained in 1/3X MMR for specified periods of regeneration time, then mounted for imaging.

Cilia length was measured in Fiji (Schindelin et al., 2012) from Z-stack images acquired of each timepoint. A segmented line was drawn along the length of each cilia with a clearly defined base and tip. Ciliary length was quantified from a minimum of 20 cells from at least 2 embryos per time-point.

Additionally, FRAP was performed on embryos at different stages of cilia regeneration. A minimum of 40 cells were imaged from at least 3 embryos per time-point.

### Analysis of IFT binding kinetics from FRAP curve fitting

For verification of binding dominant regime in FRAP experiments, the ROI was reduced from 5 μm^2^ to 1 μm^2^ in experiments and analysis, or only when analyzing as indicated. For experiments and analysis, a ROI of 1×1 μm was bleached and this same area was measured as described above in post-bleach image analysis. For analysis only, a 2.5×2.0 μm was still bleached. However, in post-bleach image analysis, a ROI focusing on the center 1×1 μm of the 2.5×2.0 μm bleached region was measured.

Fluorescence recovery as a function of time was fit to the following equation: *f*(*t*) = Λ(1 – *e^-k_off_t^*) utilizing custom nonlinear model fitting within Matlab’s curve fitting toolbox.

### Monte-Carlo model of diffusion to capture

Diffusion and binding in a 3D-environment were modeled as a random walk process, with simulations run in Matlab. 20 thousand randomly distributed particles move in a 2×2×4 μm cubic lattice with reflecting boundary conditions. If the molecules move within an apical capture radius of 400×400 nm, simulating the basal body, they have the probability of binding, dependent on their binding kinetics (*k*on*). Unbinding from the basal body is stochastic, but with a probability influenced by *k*_off_.

To ensure proper simulation of diffusion in the model, the cubic lattice was expanded (to prevent confined diffusion), *k*on* and *k*_off_ were set to zero. In this scenario, we saw appropriate scaling of area explored, with < Δ*x*^2^(*t*) > = 6*Dt*. Additionally, proper binding was determined by ensuring the number of molecules bound, *pBound*, scaled linearly with increases in *k*on*/*k*_off_.

### Statistical analysis and data visualization

Plots were generated using custom R scripts, utilizing the gglplot2 package (Wickham, 2016). One-way analysis of variance (ANOVA) comparison of FRAP mobile fraction values between basal-body proteins and of ciliary length measurements during regeneration were carried out utilizing the afex R package (Singmann et al., 2015). Differences between treated and non-treated mobile fraction values were analyzed using Welch’s t-test. P-values greater than 0.05 are reported as non-significant (“n.s”), 0.05>p>0.005 (“*”), 0.005>p>0.0005 (“**”), and p<0.005 (“***”).

## SUPPLEMENTAL MATERIALS

**Supplemental Figure 1.**
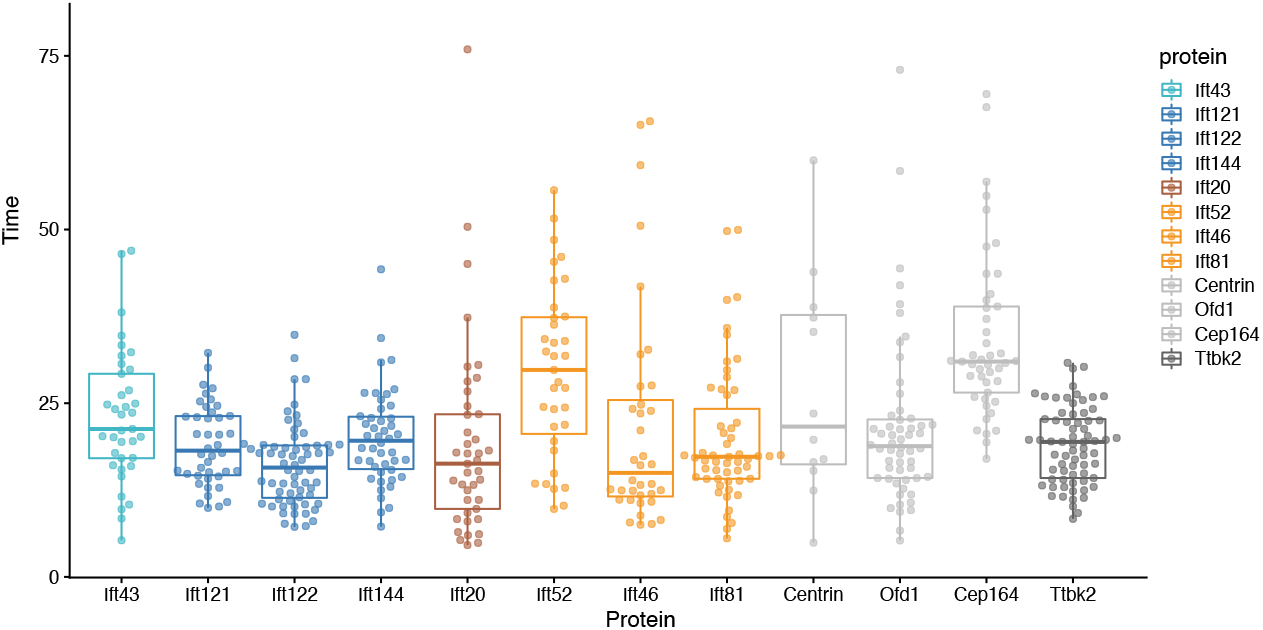
Half-times of basal body proteins are generally short. IFT and other basal body proteins generally display half-times less than 25 seconds.

**Supplemental Figure 2.**
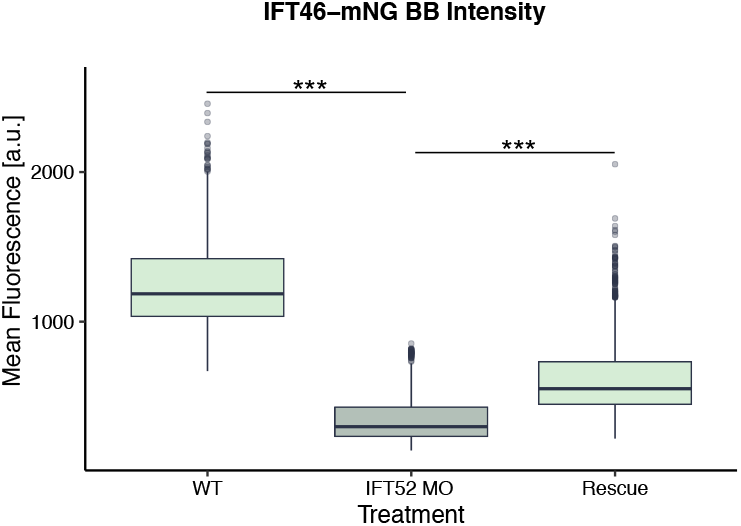
Morpholino-based KD of *ift52* is rescued by addition of Ift52. Basal body intensity of Ift46-mNG was measured in WT conditions and upon *ift52* KD and *ift52* KD + rescue.

**Supplemental Table 1.** FRAP measurements for all basal body proteins.

| Protein | [Mobile Fraction] | [Half-time] | n (cells) | n (embryos) |
| --- | --- | --- | --- | --- |
| Ift121-GFP | 56.9 | 19.0 | 42 | 5 |
| GFP-Ift122 | 57.3 | 16.1 | 64 | 12 |
| Ift144-GFP | 56.6 | 20.0 | 46 | 9 |
| Ift43-GFP | 47.7 | 23.1 | 37 | 5 |
| Ift20-GFP | 36.9 | 20.1 | 41 | 9 |
| Ift46-mNG | 46.9 | 25.7 | 36 | 10 |
| Ift52-mNG | 47.1 | 21.0 | 37 | 5 |
| Ift81-GFP | 43.4 | 20.1 | 55 | 11 |
| Centrin2-BFP | 15.6 | 27.1 | 12 | 3 |
| Cep164-GFP | 11.5 | 34.2 | 45 | 6 |
| GFP-Ofd1 | 22.6 | 21.4 | 56 | 7 |
| GFP-Ttbk2 | 64.3 | 19.0 | 70 | 9 |
| GFP-Ccdc66 | 64.9 | 55.8 | 41 | 8 |

**Supplemental Table 2.** Full statistical analysis of basal body mobile fraction comparisons.

|  | Ift121 | Ift122 | Ift144 | Ift43 | Ift20 | Ift46 | Ift52 | Ift81 | Centrin2 | Cep164 | Ofd1 | Ttbk2 |
| --- | --- | --- | --- | --- | --- | --- | --- | --- | --- | --- | --- | --- |
| Ift121 | x |  |  |  |  |  |  |  |  |  |  |  |
| Ift122 | 1.0000 | x |  |  |  |  |  |  |  |  |  |  |
| Ift144 | 1.0000 | 1.0000 | x |  |  |  |  |  |  |  |  |  |
| Ift43 | 0.0007 | <.0001 | 0.0009 | x |  |  |  |  |  |  |  |  |
| Ift20 | <.0001 | <.0001 | <.0001 | <.0001 | x |  |  |  |  |  |  |  |
| Ift46 | 0.0001 | 0.0002 | 0.0002 | 1.0000 | 0.0001 | x |  |  |  |  |  |  |
| Ift52 | 0.0003 | 0.0004 | 0.0004 | 1.0000 | 0.0002 | 1.0000 | x |  |  |  |  |  |
| Ift81 | <.0001 | <.0001 | <.0001 | 0.6058 | 0.0389 | 0.8609 | 0.8096 | x |  |  |  |  |
| Centrin2 | <.0001 | <.0001 | <.0001 | <.0001 | <.0001 | <.0001 | <.0001 | <.0001 | x |  |  |  |
| Cep164 | <.0001 | <.0001 | <.0001 | <.0001 | <.0001 | <.0001 | <.0001 | <.0001 | 0.5409 | x |  |  |
| Ofd1 | <.0001 | 0.0000 | <.0001 | <.0001 | <.0001 | <.0001 | <.0001 | <.0001 | 0.9813 | <.0001 | x |  |
| Ttbk2 | 0.0029 | 0.0009 | 0.0009 | <.0001 | <.0001 | <.0001 | <.0001 | <.0001 | <.0001 | <.0001 | <.0001 | x |
| Ccdc66 | 0.0057 | 0.0029 | 0.0023 | <.0001 | <.0001 | <.0001 | <.0001 | <.0001 | <.0001 | <.0001 | <.0001 | 1.0000 |

